# Auranofin potentiates cisplatin response through context-dependent NOTCH-associated signaling states in endometrial cancer

**DOI:** 10.64898/2026.05.27.728338

**Authors:** Robert J Lake, Cynthia Tshibangu, Niko J Candia, Quinn U Abfalterer, Irina V Lagutina, Christine Pauken, Kimberly K Leslie, Mara P Steinkamp, Hua-Ying Fan

## Abstract

Therapeutic resistance remains a major challenge in advanced and recurrent endometrial cancer (EC). Aberrant NOTCH signaling has been associated with aggressive tumor behavior and therapeutic resistance across multiple malignancies, yet its therapeutic significance in EC remains incompletely defined. We investigated whether auranofin (AuR), a noncanonical modulator of NOTCH signaling through the transcriptional effector RBPJ, alters platinum responsiveness in EC models. Elevated NOTCH3 copy-number was associated with poorer overall survival in the TCGA-UCEC cohort. AuR treatment reduced RBPJ occupancy at canonical NOTCH target loci, including *HES1* and *HES4*, across multiple EC models. Stable NOTCH3 depletion altered AuR responsiveness in a context-dependent manner while significantly enhancing cisplatin (CDDP) sensitivity in AN3CA cells. Pharmacologic AuR treatment similarly potentiated CDDP response in AN3CA cells and AN3CA xenografts, resulting in reduced tumor burden and prolonged endpoint-free survival following combination treatment. In contrast, ARK-1 xenografts demonstrated limited additional benefit from combined AuR plus CDDP therapy despite detectable suppression of RBPJ occupancy. Together, these findings identify context-dependent NOTCH-associated therapeutic vulnerabilities in EC and support further development of biomarker-guided AuR-based platinum-sensitization strategies.

**Statement of significance:** Auranofin suppresses RBPJ-associated transcriptional activity and enhances cisplatin response in biologically distinct subsets of endometrial cancer.

## Introduction

Endometrial cancer (EC) is the most common gynecologic malignancy in the United States and remains one of the few cancers for which both incidence and mortality continue to rise despite advances in cancer diagnosis and treatment (1-3). Although carboplatin- and paclitaxel-based chemotherapy remains the standard first-line treatment for advanced and recurrent EC, many patients ultimately develop recurrent or chemotherapy-refractory disease. Therefore, identifying mechanism-based therapeutic strategies that enhance chemotherapy response remains a major unmet clinical need.

The Notch signaling pathway regulates cell proliferation, differentiation, stem-cell maintenance, and survival, and aberrant NOTCH activation contributes to tumor progression and therapeutic resistance across multiple malignancies (4-14). In EC, increasing evidence supports clinically relevant roles for NOTCH signaling, particularly NOTCH2 and NOTCH3 (15). Clinical and molecular studies have linked NOTCH pathway activation in aggressive EC states and therapeutic resistance, including studies implicating aberrant NOTCH2 signaling in recurrent endometrioid EC (16). In addition, FGFR2-mutant EC models exhibit enhanced EGFR and NOTCH signaling, further supporting functional interactions between Notch signaling and oncogenic pathways in EC (15,17-21). Although the role of NOTCH signaling has been extensively investigated in ovarian cancer (22-27), comparatively fewer studies have directly examined NOTCH-directed therapeutic strategies in EC.

While the NOTCH pathway represents an attractive therapeutic target, clinical development of canonical NOTCH inhibitors has been limited by pathway complexity and dose-limiting toxicities (28). γ-secretase inhibitors suppress NOTCH activation through inhibition of NICD processing but also affect numerous additional substrates, resulting in dose-limiting gastrointestinal and immune toxicities (29). Similarly, receptor- or ligand-directed approaches have demonstrated limited therapeutic windows in solid tumors (30). More recently, the transcription-complex inhibitor CB-103 demonstrated on-target pharmacodynamics and disease stabilization in early phase I trials (NCT03422679), but objective responses were rare, and its once-daily oral dosing yielded only transient exposures (31,32). These limitations highlight the need for alternative strategies capable of suppressing NOTCH-dependent transcriptional programs.

We previously identified auranofin (AuR), an FDA-approved gold-containing compound, using a biochemical screening platform designed to identify small molecules that disrupt RBPJ– DNA interaction, a central downstream event in canonical NOTCH signaling (33). Subsequent studies demonstrated that AuR suppresses NOTCH-associated transcriptional programs in T-cell acute lymphoblastic leukemia models and reduces RBPJ occupancy at canonical NOTCH target promoters (33). More recently, we demonstrated that AuR enhances cisplatin (CDDP) efficacy in NOTCH-dependent ovarian cancer models, including patient-derived organoids and xenografts (34). In addition to its effects on NOTCH-associated transcriptional regulation, AuR is also a known inhibitor of thioredoxin reductase and can perturb cellular redox homeostasis (35), suggesting that AuR may exert antitumor activity through simultaneous modulation of NOTCH-associated transcriptional programs and oxidative stress pathways.

In the present study, we investigated the relationship between NOTCH-associated signaling states and therapeutic response to AuR across a panel of EC models. We examined the effects of AuR on RBPJ chromatin occupancy, NOTCH3-dependent therapeutic response, and platinum sensitivity in vitro and in vivo. Our findings demonstrate that AuR-mediated enhancement of CDDP response is highly context dependent across EC models and support further investigation of AuR-based combination strategies in biologically defined subsets of EC.

## Materials and Methods

### Cell lines and cell culture

RL95-2 (ATCC CRL-1671, RRID:CVCL_0505), AN3CA (ATCC HTB-111, RRID:CVCL_0028), and KLE (ATCC CRL-1622, RRID:CVCL_1329) EC cell lines were cultured in RPMI 1640 medium (VWR,CAT# 16777-145) supplemented with 10% fetal bovine serum (VWR, cat # 89510-190), 1% penicillin-streptomycin (VWR, cat# 16777-164), and 2 mmol/L L-glutamine (Gibco, cat# 25030081) at 37°C in a humidified atmosphere containing 5% CO2. ARK-1 and ARK-2 cells were obtained from Dr.Alessandro D. Santin, whereas AN3CA, RL95-2, and KLE cells were obtained from Dr. Kimberly K. Leslie. Cell lines were routinely monitored for morphology and growth characteristics and were periodically tested for mycoplasma contamination using the InvivoGen MycoStrip Mycoplasma Detection Kit. Short tandem repeat authentication was performed by ATCC (Cat# 135-XV-10).

### Protein quantification and immunoblotting

Cells were lysed in 1× SDS sample buffer containing 2.5% SDS, 125 mmol/L Tris-HCl (pH 6.8), and 20% glycerol, without bromophenol blue or dithiothreitol. Lysates were sonicated using a Branson Sonifier 101-135-126 at 25% amplitude for 60 seconds. Protein concentrations were quantified using the Pierce BCA Protein Assay Kit (Thermo Fisher Scientific, Cat# 23225). Dithiothreitol and bromophenol blue were subsequently added to the lysates. Equal amounts of protein (10–20 µg per lane) were resolved on 4%–12% Bis-Tris SurePAGE gels in 1x MOPS SDS running buffer (GenScript, Cat# M00653). Immunoblots were developed using either SuperSignal West Pico PLUS Chemiluminescent Substrate (Thermo Fisher Scientific, Cat# 34083) or WesternBright Quantum HRP substrate (Advansta, Cat# K-12042-D10) and imaged using a Konica Processor SRX-101A or a LI-COR Odyssey Fc Imaging System.

### Antibodies

Rabbit polyclonal anti-NOTCH3 antibody was from ABclonal Technology (Cat# A13522, RRID:AB_2760384; 1:2,000). Monoclonal anti-GAPDH antibody was from EMD Millipore (Cat# MAB374, RRID:AB_2107445; 1:10,000). HRP-conjugated secondary antibodies from Cell Signaling Technology included goat anti-rabbit IgG (Cat# 7074, RRID:AB_2099233; 1:3,000) and horse anti-mouse IgG (Cat# 7076, RRID:AB_330924; 1:6,000).

### Chromatin immunoprecipitation (ChIP)

ChIP assays were performed as previously described (33,34,36,37). Briefly, approximately 1 × 10^7^ cells were fixed with 1% formaldehyde for 10 minutes and sonicated on ice at 40% amplitude (30 seconds on, 90 seconds off, for 24 minutes) using a Branson Sonifier 101-135-126. Immunoprecipitation was performed using a polyclonal anti-RBPJ antibody (FC31;1:100) (38) and blocked Protein A agarose beads (Thermo Fisher Scientific, Cat# 20333). Samples were reverse cross-linked at 65°C for 16 hours. Purified ChIP DNA was analyzed by quantitative real-time PCR using PerfeCTa SYBR Green FastMix, Low ROX (Quantabio) in 384-well format on a QuantStudio 5 Real-Time PCR System (Applied Biosystems). Enrichment of RBPJ at specific genomic regions was normalized to the corresponding input DNA using the ΔCt method (2^Ct−Ctinput^). Primer sequences were shown in Table S1.

### Lentiviral shRNA-mediated NOTCH3 knockdown

NOTCH3-targeting short hairpin RNAs (shRNAs; GeneCopoeia, Cat# HSH011876-LVRU6GP) and a scrambled control shRNA (GeneCopoeia, Cat# CSHCTR001-LVRU6GP) were used for gene silencing studies. Lentiviral particles were generated using third-generation packaging plasmids as previously described (34). Cells were infected at approximately 20% confluency, and the medium was replaced after 24 hours. Seventy-two hours after infection, cells were harvested for Western blot analysis of NOTCH3 expression. To establish stable knockdown cell lines, infected cells were selected with puromycin (2 µg/mL) for 7–10 days. Stable suppression of NOTCH3 was verified by Western blotting.

### Drug treatment and cell viability measurement

For drug treatment experiments, 25,000–50,000 cells were seeded into 96-well plates and treated with varying concentrations of cisplatin (CDDP, MedChemExpress, Cat# HY-17394) and auranofin (AuR; Cayman Chemical, Cat# 15316). After 72 hours of treatment, cell viability was measured using the ATP-Glo Bioluminometric Cell Viability Assay (Biotium, Cat# 30020). Luminescence signals were measured using a BioTek Synergy Neo2 plate reader. Selected experiments were independently repeated using crystal violet staining–based viability assays for validation.

Relative viability values were normalized to vehicle-treated controls within each experiment. Dose-response curves were analyzed using nonlinear regression with a four-parameter logistic model in GraphPad Prism software (RRID:SCR_002798). IC50 values were calculated from independent biological replicates. In experiments in which dose-response curves exhibited nonparallel behavior and converged at higher drug concentrations, overall drug response was quantified using area-under-the-curve (AUC) analysis.

### Xenograft studies

All mouse procedures were approved by the University of New Mexico Animal Care and Use Committee (protocol no. 25-201709-HSC), in accordance with NIH guidelines for the care and use of experimental animals. For AN3CA xenograft studies, cells were implanted subcutaneously into NOD/SCID gamma (NSG) immunocompromised mice (RRID: IMSR_JAX:00555). Treatment was initiated three days following implantation. Mice received vehicle, AuR (6 mg/kg, intraperitoneally, 5 times weekly), CDDP (1.25 mg/kg, intraperitoneally, once weekly), or combined AuR plus CDDP treatment for 25 days.

For ARK-1 xenograft studies, cells were also implanted subcutaneously into NSG mice. In Study 2, treatment was initiated 17 days following implantation to model established tumors. In Study 1, treatment was initiated three days following implantation. Drug dosing schedules matched those used in AN3CA studies.

Tumor dimensions were measured twice weekly using calipers, and tumor volumes were calculated using the formula: V=(L×W^2^)/2, where (L) represents tumor length and (W) represents tumor width. Humane endpoint criteria included tumor volume ≥ 2000 mm^3^ or tumor-associated ulceration. Additional humane endpoint criteria included body weight loss exceeding 20% of initial body weight.

### Ethics Statement

All animal studies were approved by the Institutional Animal Care and Use Committee (IACUC) at the University of New Mexico and were performed in accordance with institutional and NIH guidelines.

### Survival analysis of NOTCH3 alterations in TCGA-UCEC

Clinical and genomic data for uterine corpus endometrial carcinoma (UCEC) were obtained from TCGA via the UCSC Xena browser (https://xenabrowser.net/). Overall survival (OS) time and status were extracted from TCGA clinical annotations. Kaplan–Meier survival curves were generated using the UCSC Xena browser. Numbers at risk are indicated for each time point.

### Statistical analysis

Comparisons between two groups were performed using two-tailed unpaired Student’s t-tests. Comparisons among multiple groups were performed using one-way ANOVA with multiple-comparison correction or Kruskal–Wallis tests with Dunn’s multiple comparisons, as indicated in figure legends. Tumor growth curves were analyzed using mixed-effects models with treatment and time as fixed effects. Kaplan–Meier survival analyses were evaluated using log-rank tests. P values < 0.05 were considered statistically significant.

### Data Availability

All data generated or analyzed during this study are included in this published article and its supplementary information files. Additional source data are available from the corresponding author upon reasonable request.

## RESULTS

### NOTCH3 copy-number gain is associated with poor clinical outcome in endometrial cancer (EC)

To assess the clinical relevance of NOTCH3 alterations in EC, we analyzed overall survival in the TCGA-UCEC cohort (Fig. 1). Stratification by NOTCH3 copy-number using segment-level DNAcopy values revealed a significant association with patient outcome. Tumors with elevated NOTCH3 copy-number (top quartile) exhibited reduced overall survival compared with tumors harboring low copy-number (bottom quartile) (Fig. 1A; log-rank *P* = 0.0013). Survival separation was most pronounced during earlier phases of follow-up, consistent with an association between increased NOTCH3 dosage and adverse clinical behavior.

**Figure 1.**
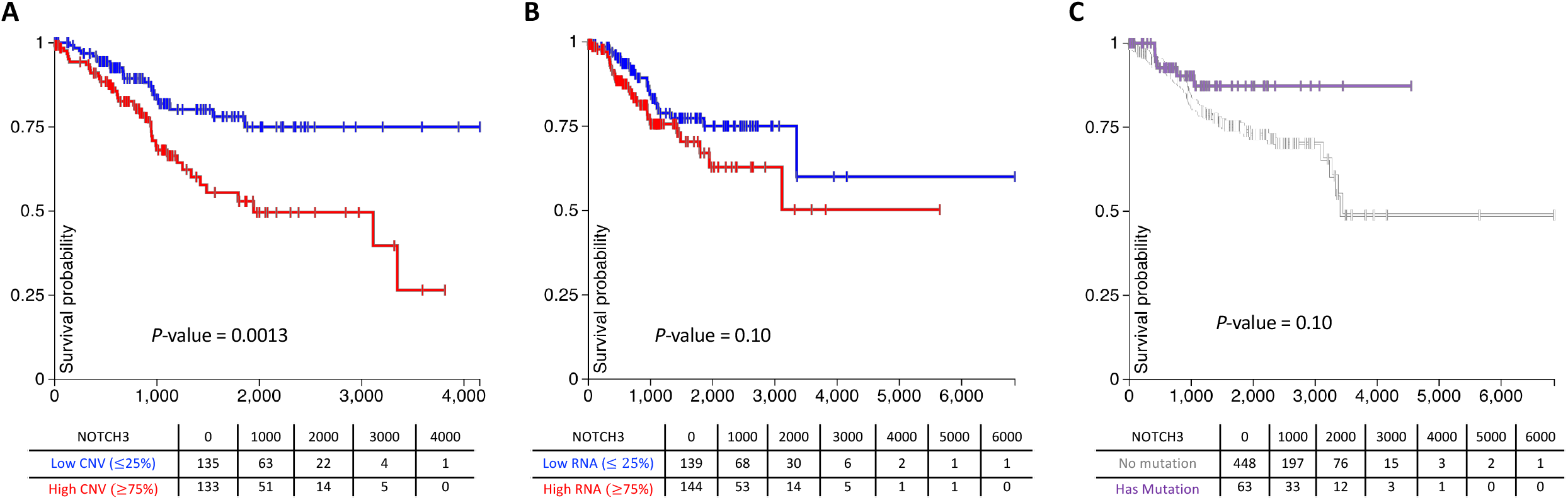
Elevated NOTCH3 copy-number is associated with poor clinical outcome in endometrial cancer. (A) Kaplan–Meier analysis of overall survival in the TCGA-UCEC cohort stratified by NOTCH3 copy-number using upper-versus-lower quartile grouping. Tumors with elevated NOTCH3 copy-number exhibited significantly reduced overall survival compared with tumors harboring low copy-number. (B) Kaplan–Meier analysis of overall survival stratified by NOTCH3 RNA expression using upper-versus-lower quartile grouping. NOTCH3 RNA expression demonstrated only a modest, nonsignificant association with outcome. (C) Kaplan–Meier analysis of overall survival according to NOTCH3 mutation status. No significant association between NOTCH3 mutation status and overall survival was observed. Log-rank *P* values are indicated on each panel. Tables beneath Kaplan–Meier plots indicate the number of patients at risk at the indicated time points (days).

To determine whether this relationship was dependent on the stratification strategy, additional robustness analyses were performed (Supplementary Fig. S1A). Binary classification of NOTCH3 copy-number status weakened survival separation relative to top-versus-bottom quartile comparisons, while continuous Cox proportional hazards modeling identified only a nonsignificant trend between NOTCH3 copy-number and overall survival (HR = 1.88, 95% CI: 0.91–3.88, *P* = 0.088). These findings suggest that the prognostic impact of NOTCH3 is most evident at the upper end of the copy-number distribution rather than through a simple linear relationship with outcome.

Using the same upper-versus-lower quartile stratification approach, NOTCH3 RNA expression demonstrated only a modest, nonsignificant association with survival (Fig. 1B), suggesting that transcript abundance alone may not adequately capture biologically relevant NOTCH3 pathway activity. Consistent with this observation, median-based stratification of NOTCH3 RNA expression similarly yielded a weak association with outcome (Supplementary Fig. S1B). Likewise, NOTCH3 mutation status was not significantly associated with overall survival (Fig. 1C), consistent with the relatively low frequency and heterogeneous functional consequences of NOTCH3 mutations in EC.

Collectively, these findings support a threshold-dependent relationship between elevated NOTCH3 copy-number and adverse clinical outcome in EC. We therefore next investigated whether NOTCH3-associated signaling contributes to therapeutic vulnerability in aggressive EC models.

### Endometrial cancer models exhibit heterogeneous NOTCH3 activity and differential therapeutic response profiles

To evaluate the relationship between NOTCH3 activity and therapeutic response in EC, we characterized a panel of EC cell lines for active NOTCH3 expression and sensitivity to auranofin and cisplatin (Fig. 2). Western blot analysis demonstrated substantial heterogeneity in NOTCH3 activation across models, as assessed by detection of the cleaved intracellular NOTCH3 fragment (NICD3) (Fig. 2A). ARK-1 and RL95-2 cells exhibited relatively high NICD3 levels, whereas AN3CA and ARK-2 displayed intermediate NICD3 expression. In contrast, KLE cells showed markedly reduced NICD3 abundance.

**Figure 2.**
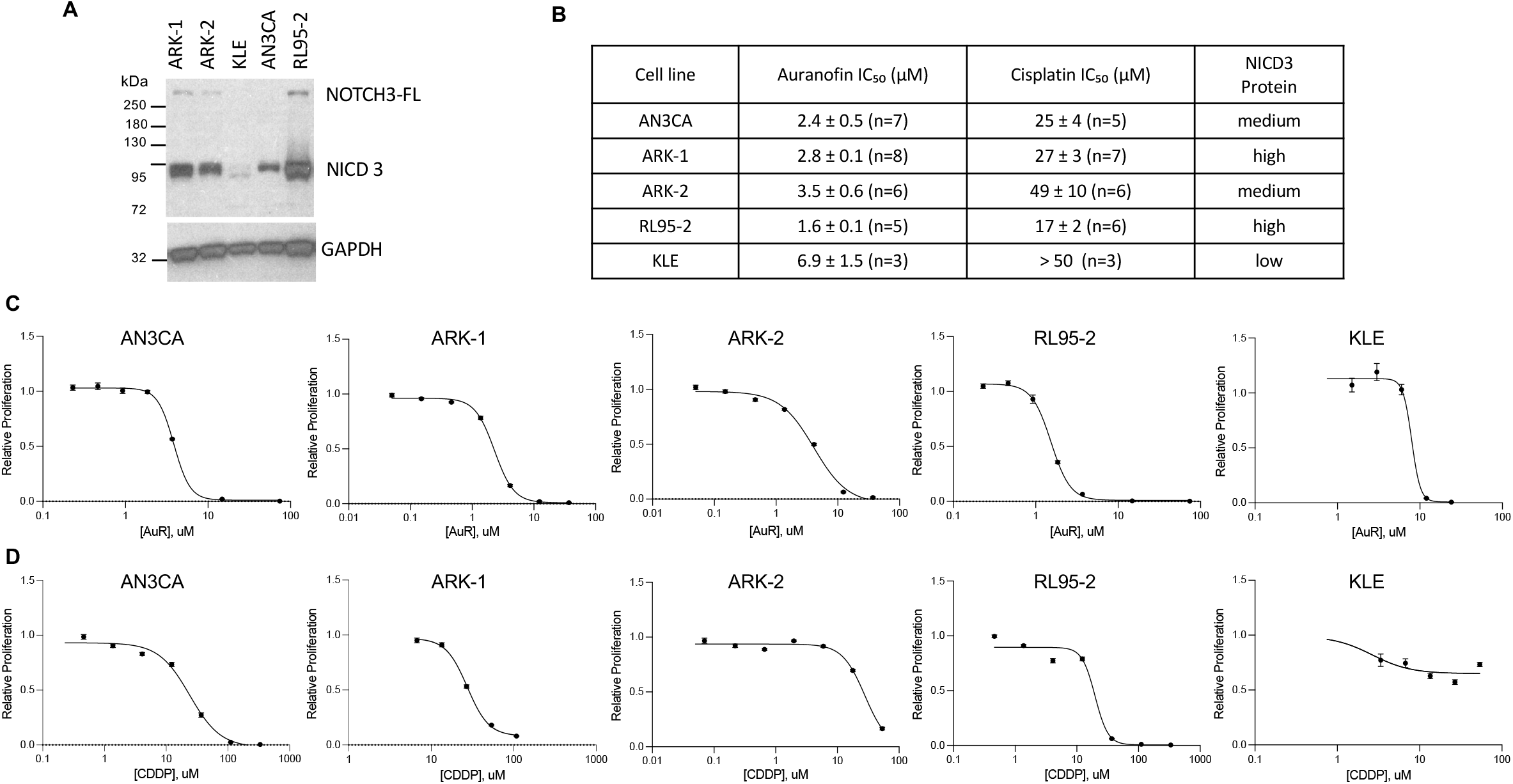
NOTCH3 protein expression and sensitivity to AuR and CDDP in EC cell lines. (A) Western blot analysis of full-length NOTCH3 (NOTCH3-FL) and activated NOTCH3 intracellular domain (NICD3) in EC cell lines. GAPDH served as a loading control. (B) Summary of AuR and CDDP IC50 values across EC cell lines. IC50 values are presented as mean ± SEM. NICD3 abundance was categorized qualitatively based on immunoblot analysis shown in panel A. (C) Dose-response analysis of AuR sensitivity in EC cell lines. (D) Dose-response analysis of CDDP sensitivity in EC cell lines. Dose-response curves in panels C and D were fitted using a four-parameter logistic regression model.

To quantitatively compare therapeutic responses across models, IC50 values for auranofin and cisplatin were determined following 72-hour treatment (Fig. 2B). RL95-2 and AN3CA cells were among the most sensitive to auranofin, whereas KLE cells exhibited relative resistance. Similarly, KLE cells displayed marked resistance to cisplatin compared with the other EC models examined. Although EC cell lines with detectable NICD3 expression generally exhibited greater sensitivity to auranofin than KLE cells, the relationship between NICD3 abundance and drug sensitivity was not strictly proportional across all models.

Representative dose-response curves for auranofin and cisplatin are shown in Fig. 2C and D, respectively. Despite broad trends between NICD3 abundance and therapeutic sensitivity, the relationship was not strictly linear across the panel. For example, ARK-2 cells exhibited intermediate NICD3 expression despite more modest auranofin sensitivity, while KLE cells combined low NICD3 abundance with relative resistance to both agents. These findings prompted us to investigate whether NOTCH3-associated signaling contributes to therapeutic vulnerability across EC models.

### AuR suppresses RBPJ occupancy and potentiates CDDP response in endometrial cancer cells

Given the differential NICD3 expression and AuR responsiveness observed across EC models (Fig. 2), we next examined whether AuR alters RBPJ occupancy at canonical NOTCH target loci and modulates platinum response. A schematic model of the proposed mechanism is shown in Fig. 3A. In RL95-2 cells, strong RBPJ enrichment above bead-only background was detected at both the *HES1* and *HES4* loci under basal conditions (Fig. 3B). Following AuR treatment, RBPJ enrichment at *HES1* was markedly reduced, while residual enrichment at *HES4* was diminished and no longer significantly above bead-only background. Similarly, ARK-1 cells demonstrated robust basal RBPJ enrichment at both *HES1* and *HES4* loci (Fig. 3C). AuR treatment reduced RBPJ occupancy at both loci. AN3CA cells also demonstrated detectable RBPJ enrichment at *HES1* and *HES4*, although the magnitude of enrichment was lower than that observed in RL95-2 and ARK-1 cells (Fig. 3D). AuR treatment reduced RBPJ occupancy at both loci.

**Figure 3.**
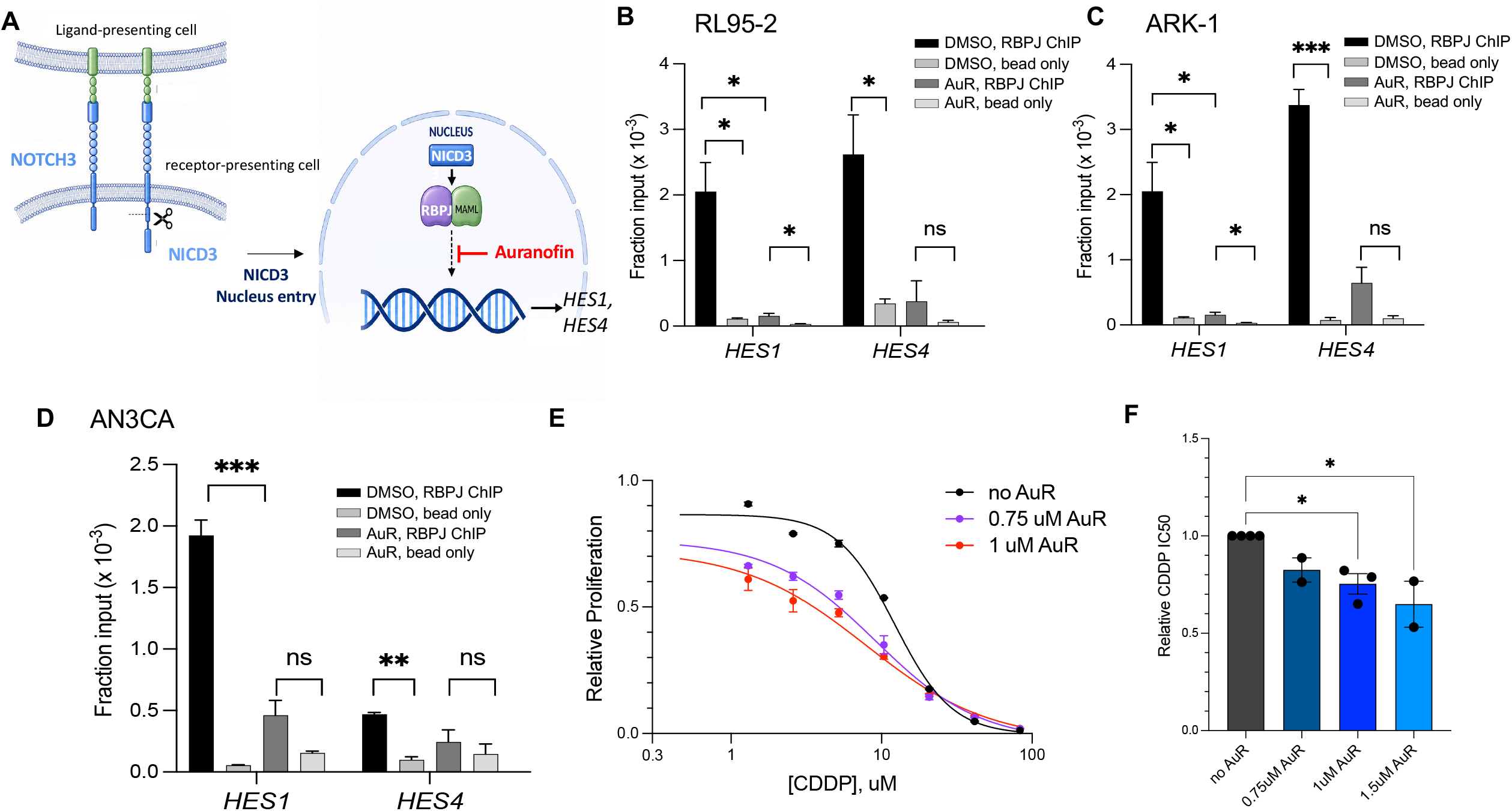
AuR suppresses RBPJ occupancy and enhances CDDP response in EC cells. (A) Schematic model illustrating canonical NOTCH3 signaling and the proposed mechanism of AuR action. Following ligand-dependent NOTCH3 activation, NICD3 forms a transcriptional activation complex with RBPJ at canonical NOTCH-responsive promoters, including *HES1* and *HES4*. AuR reduces RBPJ occupancy at canonical NOTCH-responsive promoters. (B–D) Representative RBPJ chromatin immunoprecipitation (ChIP) assays in RL95-2 (B), ARK-1 (C), and AN3CA (D) cells following treatment with DMSO or 2.5 µM AuR. RBPJ occupancy at the *HES1* and *HES4* loci was measured by qPCR and normalized to input chromatin. Bead-only controls were included to assess nonspecific background enrichment. (E) Representative nonlinear regression curves from one experiment showing CDDP response in AN3CA cells treated with increasing concentrations of CDDP in the absence or presence of 0.75 or 1.0 µM AuR. Relative proliferation was normalized to untreated controls. (F) Replicate-level quantification of relative CDDP IC50 values from independent biological replicates across AuR concentrations, including 0.75, 1.0, and 1.5 µM AuR conditions. IC50 values were normalized to the no-AuR condition. Data are presented as mean ± SEM. Statistical significance in panel F was determined using one-way ANOVA with multiple-comparison correction. ns, not significant; *, P < 0.05; **, P < 0.01; ***, P < 0.001.

To determine whether AuR alters platinum response, AN3CA cells were treated with increasing concentrations of CDDP in the presence of sub-cytotoxic concentrations of AuR. As shown in Fig. 3E, AuR treatment produced a dose-dependent leftward shift in CDDP response relative to cells treated with CDDP alone. The CDDP response curves generated in the presence of AuR maintained similar overall shapes and slopes, permitting reliable IC50 analysis. Quantification of replicate-level IC50 values demonstrated that AuR significantly reduced relative CDDP IC50 values in a dose-dependent manner (Fig. 3F).

To determine whether AuR-mediated enhancement of CDDP response extends beyond AN3CA cells, additional combination studies were performed in RL95-2 cells (Supplementary Fig. S2). AuR treatment produced a concentration-dependent leftward shift in CDDP dose-response curves. Quantification of replicate-level IC50 values demonstrated significantly reduced relative CDDP IC50 values in the presence of both 1 and 2 µM AuR. These findings further support the ability of AuR to potentiate platinum response in selected EC models.

### NOTCH3 contributes to AuR and CDDP response in a context-dependent manner

To determine whether NOTCH3 directly contributes to AuR response in EC cells, we generated stable NOTCH3 knockdown models using two independent shRNAs targeting NOTCH3 (3A and 3C) in AN3CA, KLE, and ARK-1 cells (Fig. 4A). Western blot analysis confirmed efficient depletion of NICD3 in AN3CA and ARK-1 cells, whereas KLE cells exhibited minimal baseline NICD3 expression.

**Figure 4.**
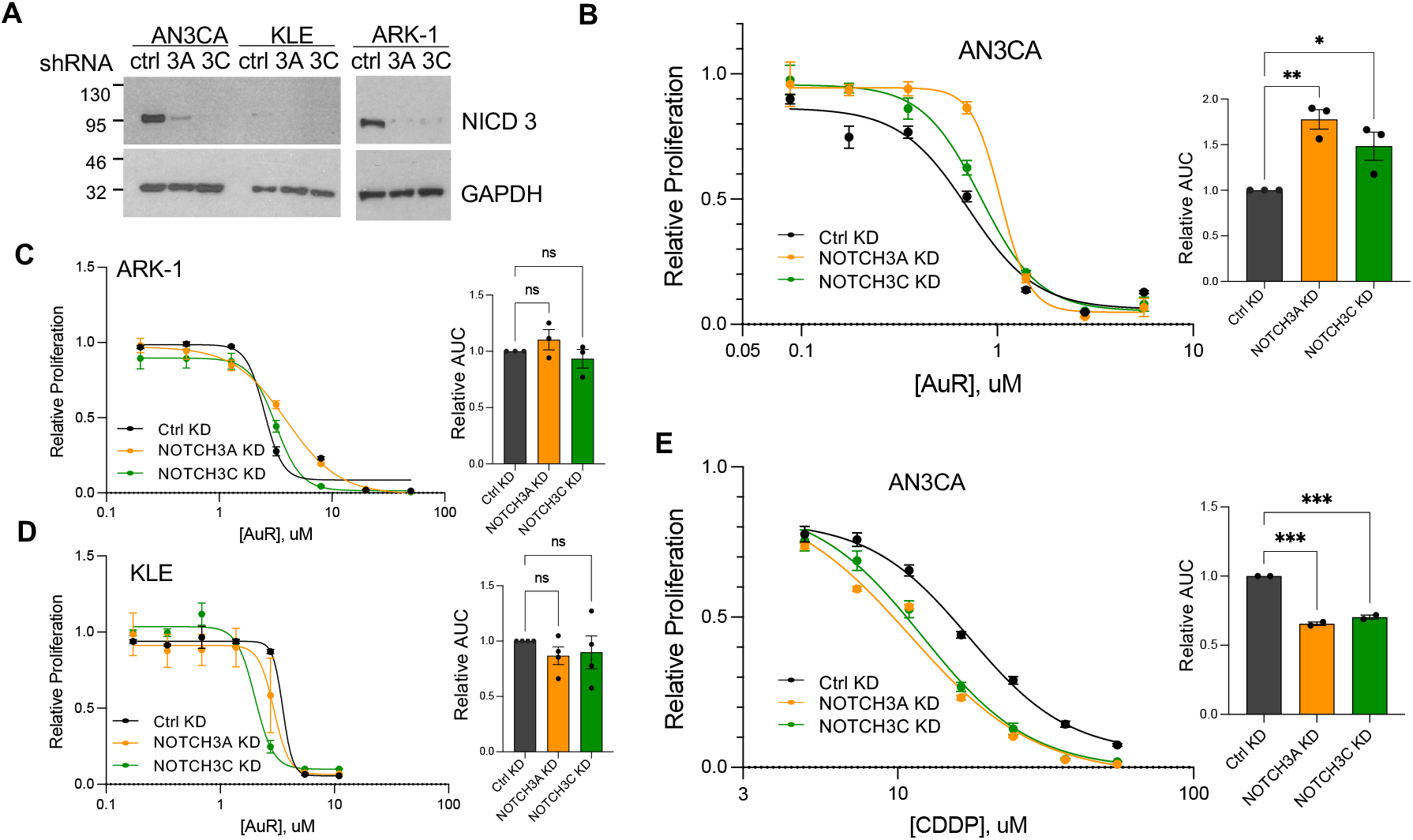
NOTCH3 depletion alters AuR and CDDP response in a context-dependent manner across EC models. (A) Western blot validation of NOTCH3 knockdown in AN3CA, KLE, and ARK-1 cells using independent shRNAs targeting NOTCH3 (3A and 3C). GAPDH served as a loading control. (B–D) Dose-response analysis of AuR response in Ctrl KD, NOTCH3A KD, and NOTCH3C KD cells. Representative nonlinear regression curves are shown at left, with replicate-level area under the curve (AUC) quantification shown at right. (E) Dose-response analysis of CDDP response in AN3CA cells following NOTCH3 depletion. Representative nonlinear regression curves are shown at left, with replicate-level AUC quantification shown at right. Relative proliferation values were normalized to vehicle-treated controls. Data are presented as mean ± SEM. AUC values were calculated from replicate dose-response curves. Statistical significance was determined using one-way ANOVA with multiple-comparison correction. *, P < 0.05; **, P < 0.01; ***, P < 0.001; ns, not significant.

We next examined the effect of NOTCH3 depletion on AuR response. In AN3CA cells, both NOTCH3A and NOTCH3C knockdown reproducibly reduced AuR sensitivity relative to control knockdown cells (Fig. 4B). NOTCH3-depleted cells maintained increased proliferation across intermediate AuR concentrations, resulting in altered dose-response behavior relative to control cells. Because the dose-response curves converged at higher AuR concentrations, IC50 values did not reliably capture these differences. Therefore, overall AuR responsiveness was quantified using area under the curve (AUC) analysis, which demonstrated significantly increased AUC values following NOTCH3 depletion. These findings indicate that NOTCH3 contributes to AuR-mediated growth suppression in AN3CA cells.

In contrast, NOTCH3 depletion produced minimal effects on AuR response in ARK-1 and KLE cells. Although knockdown modestly altered the shape of the ARK-1 dose-response curves, AUC analysis did not demonstrate a significant effect on overall AuR responsiveness (Fig. 4C). Similarly, KLE cells exhibited little change in AuR response following NOTCH3 depletion (Fig. 4D), consistent with the low baseline NICD3 expression observed in this model. Additional studies in RL95-2, ARK-1, and ARK-2 cells likewise demonstrated minimal effects of NOTCH3 depletion on therapeutic responsiveness (Supplementary Fig. S3). Together, these findings indicate that the contribution of NOTCH3 to AuR response varies substantially across EC models.

Because NOTCH3 signaling has been implicated in platinum resistance across multiple tumor types, we next examined whether NOTCH3 depletion alters CDDP response in AN3CA cells. In contrast to the modest and context-dependent effects observed with AuR, both NOTCH3A and NOTCH3C knockdown significantly enhanced CDDP responsiveness (Fig. 4E). NOTCH3-depleted cells exhibited a pronounced leftward shift in CDDP dose-response curves relative to control cells, accompanied by significantly reduced AUC values. Together, these findings demonstrate that NOTCH3 contributes to AuR response in a context-dependent manner and is associated with platinum resistance in AN3CA cells.

### AuR enhances CDDP efficacy in AN3CA xenografts

To determine whether the CDDP-potentiating effects of AuR observed *in vitro* extend to *in vivo* settings, AN3CA subcutaneous xenografts were established in NSG mice and treatment was initiated three days following tumor implantation (Fig. 5A). Mice received vehicle, AuR alone, CDDP alone, or combined AuR plus CDDP treatment for 25 days. As shown in Fig. 5B, Vehicle-treated tumors displayed continuous expansion throughout the study period. AuR monotherapy produced modest growth delay, whereas CDDP treatment partially constrained tumor progression. In contrast, combined AuR plus CDDP treatment strongly inhibited tumor expansion relative to vehicle and either monotherapy. Endpoint analysis demonstrated markedly lower tumor burden in mice receiving combination therapy compared with vehicle-, AuR-, and CDDP-treated cohorts (Fig. 5C). Consistent with these observations, Kaplan–Meier analysis using time to humane tumor endpoint demonstrated extended endpoint-free survival in mice treated with combined AuR plus CDDP relative to all other treatment groups (Fig. 5D). Collectively, these findings demonstrate that AuR potentiates CDDP antitumor activity in AN3CA xenografts.

**Figure 5.**
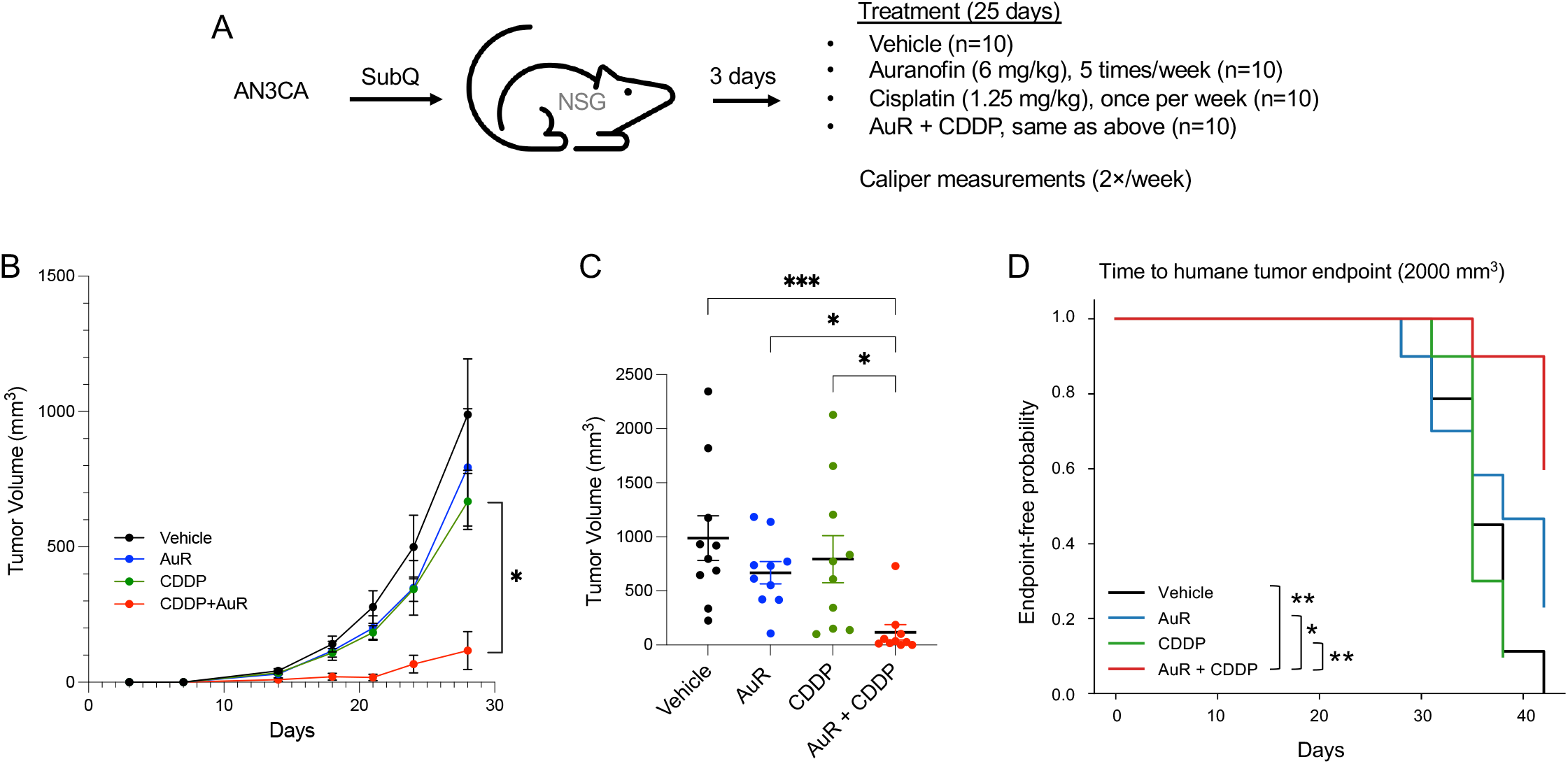
AuR enhances CDDP efficacy in AN3CA xenografts. (A) Schematic of the AN3CA subcutaneous xenograft study design. Tumor growth was monitored by caliper measurements. (B) Tumor growth curves showing mean ± SEM tumor volume over time. (C) Endpoint tumor volumes at Day 28. Each dot represents an individual mouse. (D) Kaplan–Meier analysis of time to humane tumor endpoint. Statistical significance was determined using mixed-effects model analysis (B), Kruskal–Wallis test with Dunn’s multiple-comparison correction (C), and log-rank test (D). *, P < 0.05; **, P < 0.01; ***, P < 0.001.

### AuR-mediated enhancement of CDDP response is not observed in ARK-1 xenografts

Given the high NICD3 expression and robust RBPJ occupancy observed in ARK-1 cells (Figs. 2 and 3), we next evaluated whether AuR could enhance CDDP efficacy in vivo using this model. ARK-1 cells were implanted subcutaneously into NSG mice, and treatment was initiated 17 days after implantation after tumors had become established (Fig. 6A–D).

**Figure 6.**
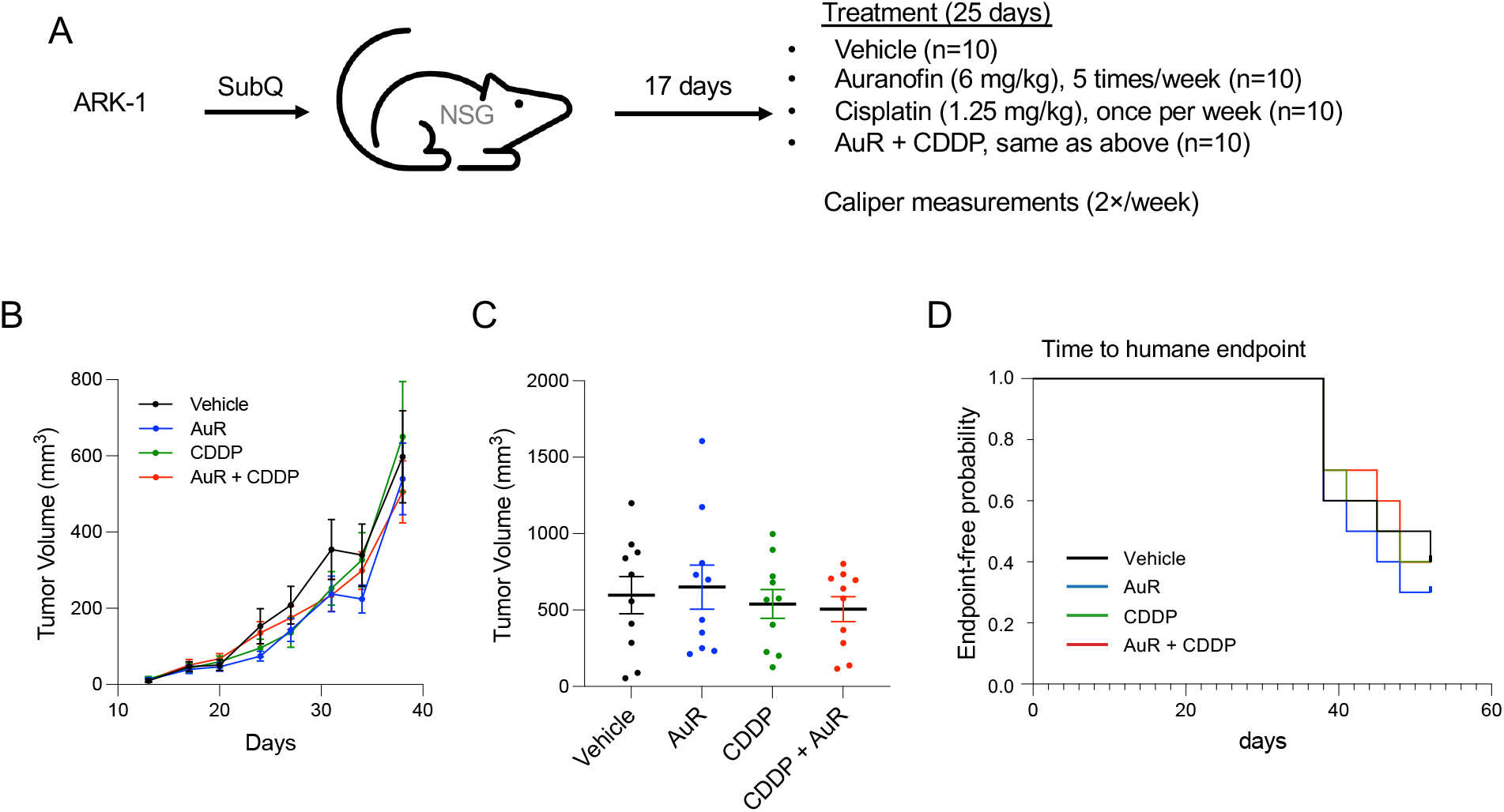
ARK-1 xenografts exhibit limited response to combined auranofin and cisplatin treatment. (A) Experimental design. ARK-1 cells were implanted subcutaneously into NSG mice. Treatment was initiated 17 days after implantation and continued for 25 days. Mice received vehicle, AuR (6 mg/kg, intraperitoneally, 5 times per week), CDDP (1.25 mg/kg, intraperitoneally, once weekly), or the combination of AuR and CDDP (n = 10 mice per group). Tumor volumes were measured twice weekly by caliper. (B) Tumor growth curves during treatment. Mean tumor volume ± SEM is shown for each treatment group. (C) Tumor volumes at day 38 (end of treatment). Each symbol represents an individual tumor; horizontal lines indicate mean ± SEM. (D) Kaplan–Meier analysis of time to humane tumor endpoint. Statistical significance was determined using mixed-effects model analysis (B), Kruskal–Wallis test with Dunn’s multiple-comparison correction (C), and log-rank test (D).

Vehicle-treated tumors progressively increased in size throughout the study (Fig. 6B). AuR monotherapy produced only modest effects on tumor growth, whereas CDDP partially suppressed tumor progression. However, combined AuR plus CDDP treatment did not further reduce tumor growth relative to CDDP alone. Consistent with these findings, endpoint tumor volumes were not significantly different between the CDDP and combination treatment groups (Fig. 6C). Similarly, Kaplan–Meier analysis revealed no clear improvement in time to humane endpoint with combined treatment compared with CDDP monotherapy (Fig. 6D).

To evaluate whether treatment timing influenced therapeutic response, an independent ARK-1 xenograft study was initiated three days after tumor implantation (Supplementary Fig. S4). Despite differences in tumor growth kinetics, combined AuR plus CDDP treatment again failed to demonstrate a reproducible improvement over CDDP alone. Together, these findings indicate that AuR-mediated enhancement of CDDP response is context dependent and may not be predicted solely by NICD3 abundance or AuR-mediated suppression of RBPJ occupancy.

## Discussion

In the present study, we demonstrate that elevated NOTCH3 copy-number is associated with adverse clinical outcome in EC (Fig. 1), that AuR suppresses RBPJ occupancy at canonical NOTCH target loci (Fig. 3), and that AuR potentiates CDDP response in selected EC models both in vitro and in vivo (Figs. 4–5, S2–S3). At the same time, our findings reveal substantial biologic heterogeneity across EC models, indicating that NOTCH-directed therapeutic vulnerabilities are highly context dependent.

Although high NOTCH3 copy-number was associated with poor prognosis, this relationship was most evident using upper-versus-lower quantile comparisons and was not maintained using continuous modeling approaches. These findings suggest that the prognostic impact of NOTCH3 may reflect threshold-dependent pathway activation rather than a simple linear relationship between pathway abundance and tumor aggressiveness. While increased NOTCH3 copy-number would generally be expected to associate with increased RNA expression, the comparatively weaker survival associations observed using bulk NOTCH3 RNA expression data suggest that transcript abundance alone may incompletely capture clinically relevant pathway-state biology. Such differences may reflect intratumoral heterogeneity, nonlinear signaling thresholds, or additional determinants of pathway activation that are not adequately represented by bulk RNA measurements.

Mechanistically, we found that AuR treatment reduced RBPJ occupancy at canonical NOTCH target loci, including *HES1* and *HES4*, across multiple EC models. These findings are consistent with prior studies identifying AuR as a noncanonical inhibitor of RBPJ-dependent transcriptional activity capable of disrupting RBPJ–chromatin interactions (33,34). At the same time, several observations indicate that canonical NOTCH3 inhibition alone does not fully explain the biologic activity of AuR. Genetic NOTCH3 depletion only partially phenocopied AuR response patterns, and AuR retained substantial cytotoxic activity in KLE cells despite minimal baseline NICD3 expression. In addition, classical NOTCH-pathway inhibitors, including the selective γ-secretase inhibitor crenigacestat and CB-103 (31,32), have not demonstrated comparable chemotherapy-potentiating activity in our endometrial cancer models. Collectively, these findings support the hypothesis that AuR exerts both NOTCH-associated and NOTCH-independent effects in EC cells.

One potential explanation for this broader activity is the well-established role of AuR as an inhibitor of thioredoxin reductase and cellular redox homeostasis (39). Because platinum agents induce oxidative and metabolic stress during chemotherapy exposure, simultaneous disruption of redox adaptation pathways could cooperate with suppression of RBPJ-dependent transcriptional programs to lower the threshold for chemotherapy-induced cytotoxicity. Such dual targeting may distinguish AuR from pathway-restricted canonical NOTCH inhibitors. Emerging evidence further supports functional crosstalk between RBPJ-associated transcriptional states and oxidative stress pathways. Previous studies have demonstrated that NOTCH/RBPJ signaling can regulate NRF2-associated cytoprotective responses and cellular antioxidant adaptation (40), while ROS-dependent NRF2–NOTCH interactions contribute to adaptive cellular signaling states (41). Consistent with these observations, AuR has also been reported to modulate NRF2/Keap1-associated stress adaptation pathways during chemotherapy sensitization (42). These observations are consistent with our finding that suppression of RBPJ promoter occupancy alone did not uniformly predict AuR responsiveness across EC models. Together, these data support the possibility that AuR response reflects integrated NOTCH/redox signaling states rather than isolated inhibition of canonical NOTCH target genes.

Importantly, AuR monotherapy produced only modest antitumor activity in most settings (Figs. 5–6), whereas substantially greater effects were observed when AuR was combined with CDDP (Fig. 5). Moreover, potentiation of platinum response was achieved using sub-cytotoxic AuR concentrations in vitro, raising the possibility that lower-dose combination strategies could provide therapeutic benefit while potentially minimizing toxicities associated with more extensive pathway inhibition. This concept may be particularly relevant given the dose-limiting gastrointestinal toxicities historically associated with pan-NOTCH inhibition using γ-secretase inhibitors.

The relationship between NOTCH3 activity and platinum response was particularly notable in AN3CA models. AN3CA is a poorly differentiated, aggressive EC model with intermediate platinum responsiveness. Genetic suppression of NOTCH3 significantly enhanced CDDP responsiveness, and pharmacologic AuR treatment similarly potentiated CDDP activity both in vitro and in vivo (Figs. 4–5). Importantly, the enhancement observed with AuR was not solely attributable to baseline cytotoxicity, as combination treatment additionally shifted CDDP dose-response relationships toward lower cisplatin concentrations. These findings are consistent with prior studies linking NOTCH signaling to platinum resistance and further support the concept that modulation of NOTCH-associated transcriptional programs can alter chemotherapy response thresholds in selected EC contexts (23,25,34,43-46). Although reduced RBPJ occupancy supports disruption of canonical NOTCH-associated chromatin engagement, future studies will be required to define the broader transcriptional consequences of AuR treatment in EC cells.

In contrast, ARK-1 xenografts demonstrated limited benefit from combined AuR plus CDDP treatment despite high NICD3 expression and detectable suppression of RBPJ occupancy following AuR exposure. An independent ARK-1 study initiated shortly after tumor implantation similarly failed to demonstrate a reproducible enhancement of CDDP efficacy by AuR. Together with the variable effects of NOTCH3 depletion observed across EC cell models, these findings highlight the context-dependent nature of AuR responsiveness. Such heterogeneity likely reflects differences in signaling dependencies among EC tumors and suggests that successful deployment of AuR-based therapeutic strategies may require biologically informed patient stratification rather than reliance on a single pathway biomarker.

Several limitations of the present study should be acknowledged. Although our findings support a role for NOTCH-associated signaling in modulating platinum response, the precise molecular mechanisms linking AuR treatment to altered chemotherapy sensitivity remain incompletely defined. In addition, the current study focused primarily on established cell line and xenograft systems, and future studies using patient-derived models will be important for defining clinically relevant biomarkers and response states. Further investigation will also be required to dissect the relative contributions of Notch-associated and redox-associated mechanisms to therapeutic response.

In summary, our findings demonstrate that AuR suppresses RBPJ occupancy at canonical Notch target loci and potentiates platinum response in selected endometrial cancer contexts. Collectively, our findings are consistent with a model in which AuR alters chemotherapy responsiveness through convergent effects on NOTCH-associated and redox-associated survival pathways and warrant continued investigation of AuR-based combination strategies in biomarker-defined subsets of endometrial cancer.

## Acknowledgements

We thank the Animal Models and Flow Cytometry Shared Resources at the UNM Comprehensive Cancer Center (UNMCCC) for their support. This work was supported by the UNMCCC Support Grant (P30CA118100; RL, IL, KL, MS, and HF), NIH grants R21CA286210 (RL, KL, MS, and HF) and R01CA99908-22 (KL), Department of Defense awards CDMRP OC230061 (RL, KL, MP, and HF) and CA210610 (KL), and the Route 66 Endometrial Cancer SPORE (P50CA265793). CT is supported by the American Cancer Society Diversity in Cancer Research (DICR) Post-Baccalaureate Fellows Program. NJC is supported by the NIH U-RISE program.

## Authors’ Contributions

RJL and HYF: Conceptualization, resources, methodology, investigation, formal analysis, validation, visualization, supervision, funding acquisition, data curation, and writing–original draft, review, and editing. CT, NJC, QUA, and CP: Data curation, formal analysis, and writing– review and editing. IL: Conceptualization, investigation, methodology, formal analysis, data curation, and writing–review and editing. KKL: Resources and writing–review and editing. MPS: Resources, supervision, methodology, and writing–review and editing.

## REFERENCES

1. Siegel RL, Kratzer TB, Giaquinto AN, Sung H, Jemal A. Cancer statistics, 2025. CA Cancer J Clin 2025;75:10–45

2. National Cancer I. SEER Cancer Stat Facts: Uterine Cancer (Corpus Uteri). 2025;https://seer.cancer.gov/statfacts/html/corp.html

3. American Cancer S. Cancer Facts &Figures 2025. https://www.cancer.org/content/dam/cancer-org/research/cancer-facts-and-statistics/annual-cancer-facts-and-figures/2025/2025-cancer-facts-and-figures-acs.pdf: American Cancer Society;2025 2025/09/29.

4. Previs RA, Coleman RL, Harris AL, Sood AK. Molecular pathways: translational and therapeutic implications of the Notch signaling pathway in cancer. Clin Cancer Res 2015;21:955–61

5. Bray SJ, Bigas A. Modes of Notch signalling in development and disease. Nature reviews Molecular cell biology 2025;26:522–37

6. Chen C, Du Y, Nie R, Wang S, Wang H, Li P. Notch signaling in cancers: mechanism and potential therapy. Front Cell Dev Biol 2025;13:1542967

7. Ghosh A, Mitra AK. Metastasis and cancer associated fibroblasts: taking it up a NOTCH. Front Cell Dev Biol 2023;11:1277076

8. Zhou B, Lin W, Long Y, Yang Y, Zhang H, Wu K, et al. Notch signaling pathway: architecture, disease, and therapeutics. Signal Transduct Target Ther 2022;7:95

9. You WK, Schuetz TJ, Lee SH. Targeting the DLL/Notch Signaling Pathway in Cancer: Challenges and Advances in Clinical Development. Mol Cancer Ther 2023;22:3–11

10. Zhdanovskaya N, Firrincieli M, Lazzari S, Pace E, Scribani Rossi P, Felli MP, et al. Targeting Notch to Maximize Chemotherapeutic Benefits: Rationale, Advanced Strategies, and Future Perspectives. Cancers (Basel) 2021;13

11. Muthuraman N, Thomas A, Samuel Ram T, Mohankumar KM, Abraham P. Is There Any Difference in Stem Cell Population between Type I and Type II Endometrial Cancer? A Pilot Study. J Mother Child 2025;29:10–9

12. Xiu M, Wang Y, Li B, Wang X, Xiao F, Chen S, et al. The Role of Notch3 Signaling in Cancer Stemness and Chemoresistance: Molecular Mechanisms and Targeting Strategies. Front Mol Biosci 2021;8:694141

13. Polychronidou G, Kotoula V, Manousou K, Kostopoulos I, Karayannopoulou G, Vrettou E, et al. Mismatch repair deficiency and aberrations in the Notch and Hedgehog pathways are of prognostic value in patients with endometrial cancer. PLoS One 2018;13:e0208221

14. Cho S, Lu M, He X, Ee PL, Bhat U, Schneider E, et al. Notch1 regulates the expression of the multidrug resistance gene ABCC1/MRP1 in cultured cancer cells. Proceedings of the National Academy of Sciences of the United States of America 2011;108:20778–83

15. Cancer Genome Atlas Research N, Kandoth C, Schultz N, Cherniack AD, Akbani R, Liu Y, et al. Integrated genomic characterization of endometrial carcinoma. Nature 2013;497:67–73

16. Devor EJ, Miecznikowski J, Schickling BM, Gonzalez-Bosquet J, Lankes HA, Thaker P, et al. Dysregulation of miR-181c expression influences recurrence of endometrial endometrioid adenocarcinoma by modulating NOTCH2 expression: An NRG Oncology/Gynecologic Oncology Group study. Gynecol Oncol 2017;147:648–53

17. Dixit G, Gonzalez-Bosquet J, Skurski J, Devor EJ, Dickerson EB, Nothnick WB, et al. FGFR2 mutations promote endometrial cancer progression through dual engagement of EGFR and Notch signalling pathways. Clin Transl Med 2023;13:e1223

18. Byron SA, Gartside M, Powell MA, Wellens CL, Gao F, Mutch DG, et al. FGFR2 point mutations in 466 endometrioid endometrial tumors: relationship with MSI, KRAS, PIK3CA, CTNNB1 mutations and clinicopathological features. PLoS One 2012;7:e30801

19. Dutt A, Salvesen HB, Chen TH, Ramos AH, Onofrio RC, Hatton C, et al. Drug-sensitive FGFR2 mutations in endometrial carcinoma. Proceedings of the National Academy of Sciences of the United States of America 2008;105:8713–7

20. Pollock PM, Gartside MG, Dejeza LC, Powell MA, Mallon MA, Davies H, et al. Frequent activating FGFR2 mutations in endometrial carcinomas parallel germline mutations associated with craniosynostosis and skeletal dysplasia syndromes. Oncogene 2007;26:7158–62

21. Krakstad C, Birkeland E, Seidel D, Kusonmano K, Petersen K, Mjos S, et al. High-throughput mutation profiling of primary and metastatic endometrial cancers identifies KRAS, FGFR2 and PIK3CA to be frequently mutated. PLoS One 2012;7:e52795

22. Hu W, Liu T, Ivan C, Sun Y, Huang J, Mangala LS, et al. Notch3 pathway alterations in ovarian cancer. Cancer Res 2014;74:3282–93

23. McAuliffe SM, Morgan SL, Wyant GA, Tran LT, Muto KW, Chen YS, et al. Targeting Notch, a key pathway for ovarian cancer stem cells, sensitizes tumors to platinum therapy. Proceedings of the National Academy of Sciences of the United States of America 2012;109:E2939–48

24. Bhattacharya R, Ghosh A, Mukhopadhyay S. High-grade serous ovarian carcinoma, the “Achiles’hill”for clinicians and molecular biologists: a molecular insight. Mol Biol Rep 2023;50:9511–9

25. Park JT, Chen X, Trope CG, Davidson B, Shih Ie M, Wang TL. Notch3 overexpression is related to the recurrence of ovarian cancer and confers resistance to carboplatin. Am J Pathol 2010;177:1087–94

26. Chen X, Thiaville MM, Chen L, Stoeck A, Xuan J, Gao M, et al. Defining NOTCH3 target genes in ovarian cancer. Cancer Res 2012;72:2294–303

27. Park JT, Li M, Nakayama K, Mao TL, Davidson B, Zhang Z, et al. Notch3 gene amplification in ovarian cancer. Cancer Res 2006;66:6312–8

28. Allen F, Maillard I. Therapeutic Targeting of Notch Signaling: From Cancer to Inflammatory Disorders. Front Cell Dev Biol 2021;9:649205

29. McCaw TR, Inga E, Chen H, Jaskula-Sztul R, Dudeja V, Bibb JA, et al. Gamma Secretase Inhibitors in Cancer: A Current Perspective on Clinical Performance. Oncologist 2021;26:e608–e21

30. Moore G, Annett S, McClements L, Robson T. Top Notch Targeting Strategies in Cancer: A Detailed Overview of Recent Insights and Current Perspectives. Cells 2020;9

31. Lehal R, Zaric J, Vigolo M, Urech C, Frismantas V, Zangger N, et al. Pharmacological disruption of the Notch transcription factor complex. Proceedings of the National Academy of Sciences of the United States of America 2020;117:16292–301

32. Hanna GJ, Stathis A, Lopez-Miranda E, Racca F, Quon D, Leyvraz S, et al. A Phase I Study of the Pan-Notch Inhibitor CB-103 for Patients with Advanced Adenoid Cystic Carcinoma and Other Tumors. Cancer Res Commun 2023;3:1853–61

33. Lake RJ, Haynes MK, Dreval K, Bilkis R, Sklar LA, Fan HY. A Novel Flow Cytometric Assay to Identify Inhibitors of RBPJ-DNA Interactions. SLAS Discov 2020;25:895–905

34. Lake RJ, Nikeghbal P, Lagutina IV, Leslie KK, Steinkamp MP, Fan HY. Auranofin Synergizes with Cisplatin in Reducing Tumor Burden of NOTCH-Dependent Ovarian Cancer. Cancer Res Commun 2025;5:1796–808

35. Ranbhise JS, Singh MK, Ju S, Han S, Yun HR, Kim SS, et al. The Redox Paradox: Cancer’s Double-Edged Sword for Malignancy and Therapy. Antioxidants (Basel) 2025;14

36. Dreval K, Lake RJ, Fan HY. HDAC1 negatively regulates selective mitotic chromatin binding of the Notch effector RBPJ in a KDM5A-dependent manner. Nucleic Acids Res 2019;47:4521–38

37. Lake RJ, Boetefuer EL, Tsai PF, Jeong J, Choi I, Won KJ, et al. The sequence-specific transcription factor c-Jun targets Cockayne syndrome protein B to regulate transcription and chromatin structure. PLoS Genet 2014;10:e1004284.

38. Lake RJ, Tsai PF, Choi I, Won KJ, Fan HY. RBPJ, the major transcriptional effector of Notch signaling, remains associated with chromatin throughout mitosis, suggesting a role in mitotic bookmarking. PLoS Genet 2014;10:e1004204

39. Abdalbari FH, Telleria CM. The gold complex auranofin: new perspectives for cancer therapy. Discov Oncol 2021;12:42

40. Wakabayashi N, Skoko JJ, Chartoumpekis DV, Kimura S, Slocum SL, Noda K, et al. Notch-Nrf2 axis: regulation of Nrf2 gene expression and cytoprotection by notch signaling. Mol Cell Biol 2014;34:653–63

41. Paul MK, Bisht B, Darmawan DO, Chiou R, Ha VL, Wallace WD, et al. Dynamic changes in intracellular ROS levels regulate airway basal stem cell homeostasis through Nrf2-dependent Notch signaling. Cell Stem Cell 2014;15:199–214

42. Deepika N, Rajendra Prasad N, Radhiga T. Auranofin sensitizes breast cancer cells to paclitaxel mediated cell death via regulating FOXO3/Nrf2/Keap1 signaling pathway. Cell Biochem Funct 2024;42:e3903

43. Liu YP, Yang CJ, Huang MS, Yeh CT, Wu AT, Lee YC, et al. Cisplatin selects for multidrug-resistant CD133+cells in lung adenocarcinoma by activating Notch signaling. Cancer Res 2013;73:406–16

44. Dai G, Deng S, Guo W, Yu L, Yang J, Zhou S, et al. Notch pathway inhibition using DAPT, a gamma-secretase inhibitor (GSI), enhances the antitumor effect of cisplatin in resistant osteosarcoma. Mol Carcinog 2019;58:3–18

45. Koucka K, Spalenkova A, Seborova K, Tesarova T, Ehrlichova M, Krus I, et al. Molecular impact of NOTCH signaling dysregulation on ovarian cancer progression, chemoresistance, and taxane response. Biomed Pharmacother 2025;191:118532

46. Leung SOA, Konstantinopoulos PA. Advances in the treatment of platinum resistant epithelial ovarian cancer: an update on standard and experimental therapies. Expert Opin Investig Drugs 2021;30:695–707

