## Supplementary materials for "Auranofin potentiates cisplatin response through context-dependent NOTCH-associated signaling states in endometrial cancer"

**Supplementary Survival Analysis Methods**

To assess the robustness of survival associations, additional analyses were performed using median-based copy-number stratification and univariate Cox proportional hazards modeling with NOTCH3 copy-number treated as a continuous variable. Kaplan–Meier analyses were performed using the UCSC Xena Browser. Cox proportional hazards modeling was performed using NOTCH3 copy-number as a continuous variable. Median-based stratification analyses were used to compare outcomes between tumors with copy-number values above and below the cohort median.

**Supplementary Table S1.**

| **Table S1. Primers used in qPCR** | |
|---|---|
| hHES1_ChIP_F | 5′-CCT CCC ATT GGC TGA AAG T-3′ |
| hHES1_ChIP_R | 5′-CGG ATC CTG TGT GAT CCC TA-3′ |
| hHES4_ChIP_F | 5′-CTC AGG CCG TTT CCC TAT TT-3′ |
| hHES4_ChIP_R | 5′-CGA GGC GTG ACT GAC AGC-3′ |

### Supplementary Figures

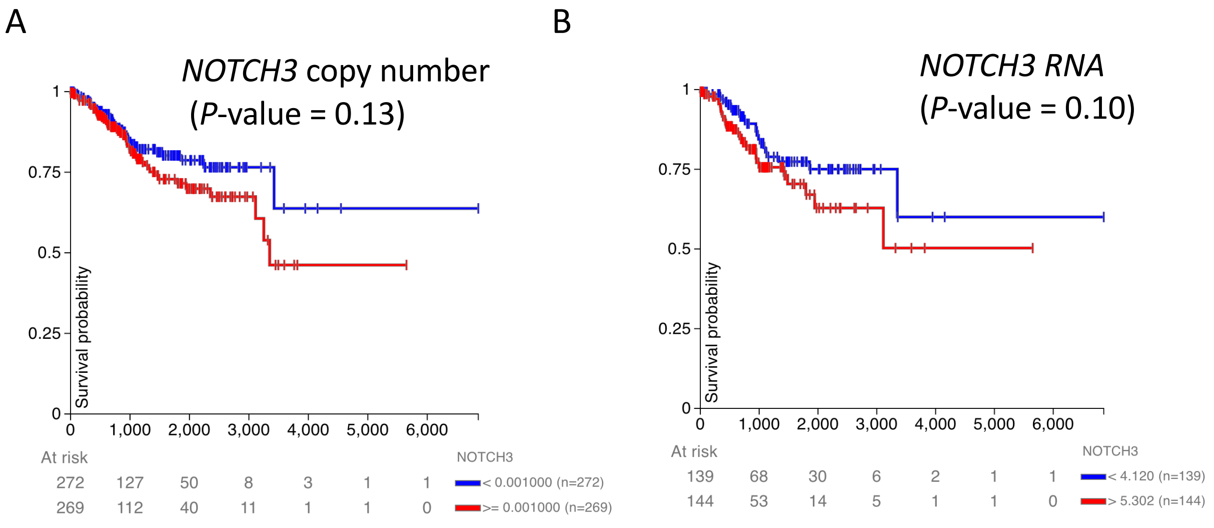


**Figure S1. Robustness analysis of NOTCH3 survival associations.**

**A,** Kaplan–Meier analysis using median-based stratification of NOTCH3 copy-number demonstrates weaker survival separation than upper-versus-lower quartile grouping, indicating that the association observed in Fig. 1A is most evident among tumors with the highest levels of copy-number gain. **B,** Kaplan–Meier analysis using median-based stratification of NOTCH3 RNA expression demonstrates a similarly modest and nonsignificant association with outcome. Together with continuous Cox proportional hazards modeling reported in the Results section, these analyses support a threshold-dependent relationship between elevated NOTCH3 copy-number and adverse clinical outcome in endometrial cancer.

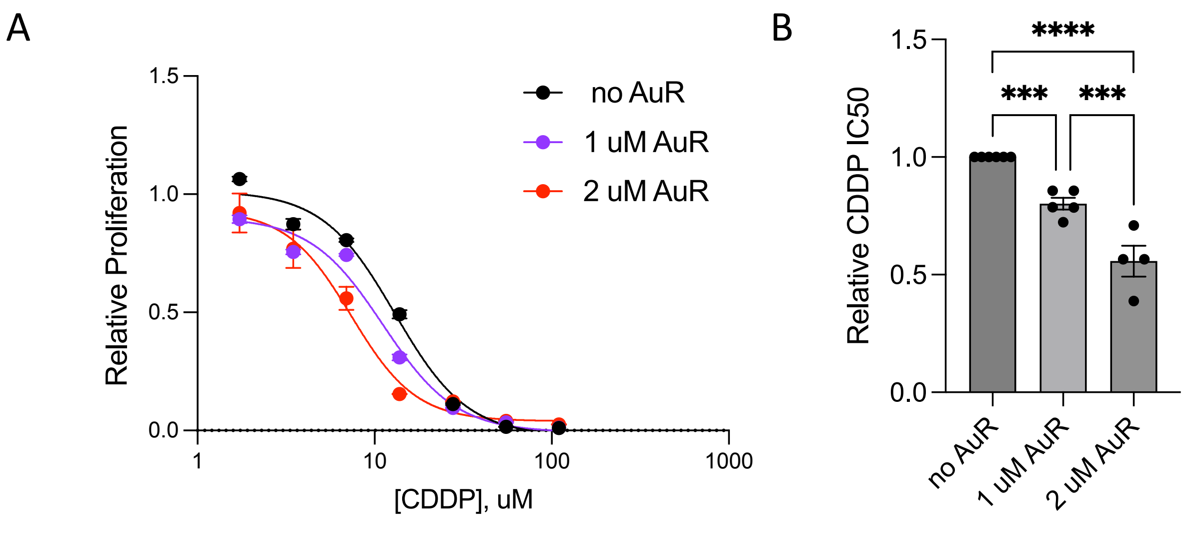


**Supplementary Figure S2. AuR enhances CDDP response in additional endometrial cancer cell models.**

RL95-2 cells were treated with increasing concentrations of CDDP in the absence or presence of 1 or 2 µM AuR. Relative proliferation was normalized to untreated controls. **A**, Representative nonlinear regression dose-response curves are shown at left. **B**, Quantification of relative CDDP IC50 values from independent biological replicates, with IC50 values normalized to the no-AuR condition. Data are presented as mean ± SEM from independent biological replicates. IC50 values were calculated from replicate dose-response curves. Statistical significance in panel B was determined using one-way ANOVA with multiple-comparison correction. ns, not significant; ***, $P < 0.001$; ****, $P < 0.0001$.

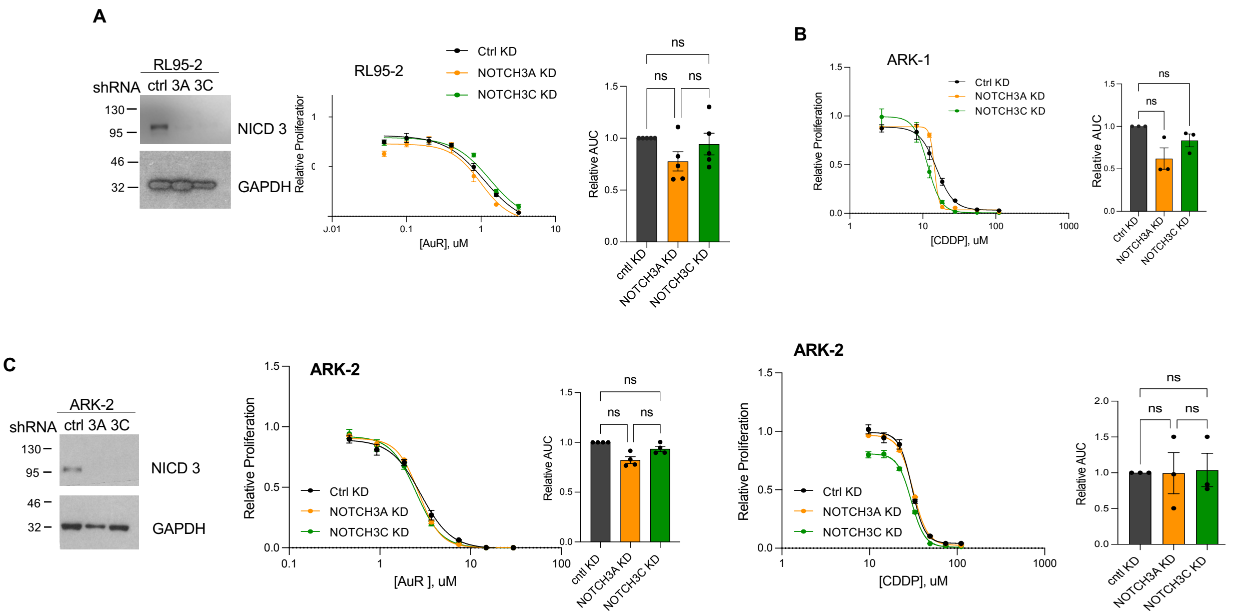


**Supplementary Figure S3. NOTCH3 depletion produces minimal effects on AuR and CDDP responsiveness in additional endometrial cancer models.**

**A**, Western blot validation of NOTCH3 knockdown and AuR dose–response analysis in RL95-2 cells. Relative proliferation curves and AUC quantification demonstrate minimal changes in AuR response following NOTCH3 depletion. **B**, CDDP dose–response analysis in ARK-1 cells following NOTCH3 knockdown. Relative proliferation curves and AUC quantification demonstrate no significant effect of NOTCH3 depletion on CDDP responsiveness. **C**, Western blot validation of NOTCH3 knockdown together with AuR and CDDP dose–response analyses in ARK-2 cells. Relative proliferation curves and corresponding AUC quantification demonstrate minimal effects of NOTCH3 depletion on either AuR or CDDP responsiveness. Data are presented as mean ± SEM from independent biological replicates. Statistical significance was determined using one-way ANOVA with multiple-comparison correction. ns, not significant.

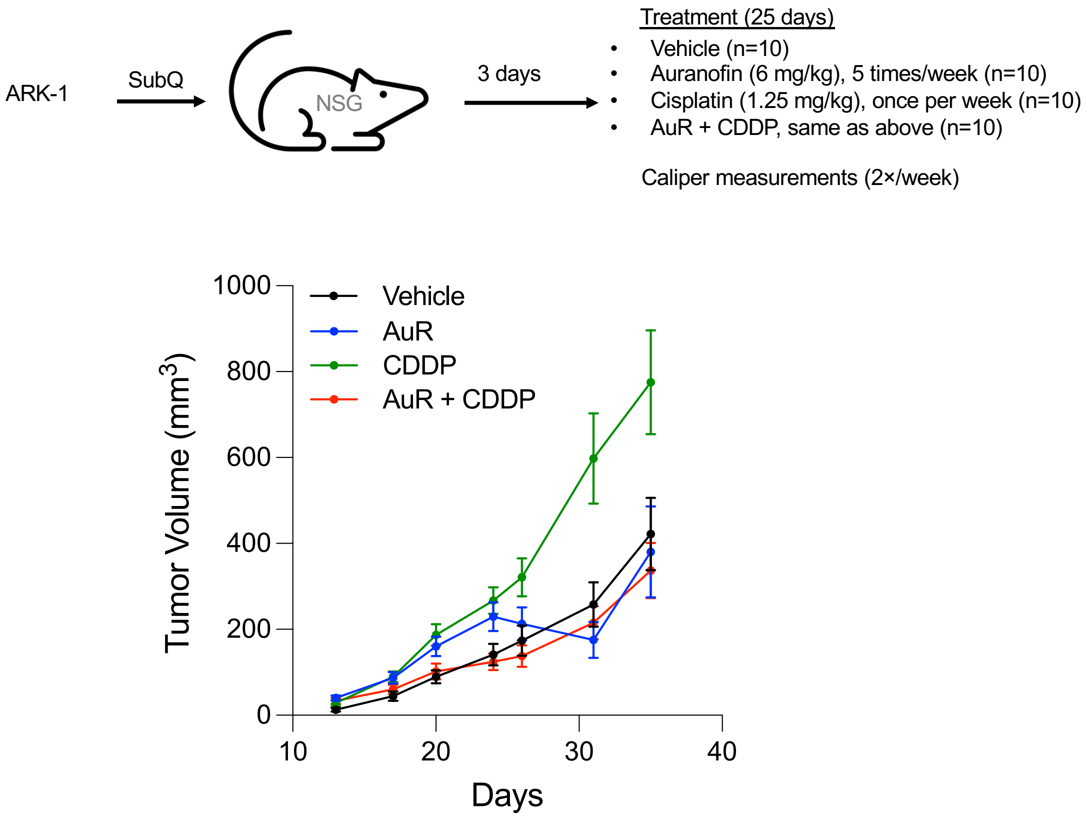


**Supplementary Figure S4. Early-treatment ARK-1 xenograft study.**

ARK-1 cells were implanted subcutaneously into NSG mice and treatment was initiated 3 days after implantation. Mice received vehicle, auranofin (AuR; 6 mg/kg, intraperitoneally, 5 times per week), cisplatin (CDDP; 1.25 mg/kg, intraperitoneally, once weekly), or the combination of AuR and CDDP (n = 10 mice per group). Tumor volumes were measured twice weekly by caliper. Tumor growth curves are shown as mean ± SEM tumor volume.
